# Downregulation of CYP7B1 caused by lipotoxicity associates with the progression of non-alcoholic steatohepatitis

**DOI:** 10.1101/2023.05.10.539979

**Authors:** Yuichi Watanabe, Kyohei Kinoshita, Napatsorn Dumrongkulchart, Takashi Sasaki, Makoto Shimizu, Yoshio Yamauchi, Ryuichiro Sato

## Abstract

Non-alcoholic fatty liver disease (NAFLD) is the most common chronic liver disease worldwide, with an incidence of >25% of the adult population. NAFLD ranges from benign simple steatosis to non-alcoholic steatohepatitis (NASH). However, its transition mechanisms underlying the pathogenesis remain to be clarified. The expression of *Cyp7b1* gene is downregulated in the liver of leptin-deficient mice and methionine and choline-deficient diet-fed mice based on previous microarray data. Thus, in this study, we investigated the effect of CYP7B1 restoration on the progression of NASH in mice fed MCD diet and its association with oxidative and lipid stresses. Our results suggest that restoration of CYP7B1 expression attenuates hepatitis and fibrosis and that lipid and oxidative stresses observed in the early stage of NASH suppresses *Cyp7b1* transcription in hepatocytes.

## Introduction

Non-alcoholic fatty liver disease (NAFLD) is the major cause of chronic liver disease affecting >25% of the world’s adult population, and its morbidity is increasing each year [1]. NAFLD ranges from relatively benign and reversible simple steatosis to non-alcoholic steatohepatitis (NASH). NASH is characterized by inflammation, hepatocellular injury, and fibrosis and steatosis [2]. It is estimated that 20% of patients with NAFLD will develop NASH [3]. NASH can progress from simple steatosis due to several factors. Insulin resistance due to fat accumulation in hepatocytes exacerbate its pathogenesis, which is simultaneously progressed by the release of inflammatory cytokines that induce inflammation and fibrosis, oxidative stress, endoplasmic reticulum stress, and abnormal mitochondrial function [2]. However, the transition mechanisms of from simple steatosis to NASH remain unclear.

CYP7B1, cytochrome P450 family 7 subfamily B member 1, is widely expressed in various tissues involved in bile acid synthesis, steroid genesis, and reabsorption. CYP7B1 in the endoplasmic reticulum catalyzes 7α-hydroxylation of 25- and 27-hydroxycholesterol (HC) into 7α,25- and 7α,27-dihydroxycholesterol, respectively [4]. Chenodeoxycholic acid is synthesized via the alternative pathway of bile acid synthesis. CYP7B1 also plays an important role in other steroid metabolisms such as hydroxylation of dehydroepiandrosterone and pregnenolone [5, 6].

From 45,101 genes that are altered in leptin-deficient (ob/ob) mice and mice fed with methionine choline-deficient (MCD) plus high-fat diet, *Cyp7b1* is one of the genes that are significantly downregulated in both models (Gene expression Omnibus; GSE49195 and GSE35961) [7, 8]. This downregulation is also observed in the liver of western diet-fed mice [9]. The inborn absence of CYP7B1 increases the markers of liver damage and giant hepatocytes, as well as subsequent cirrhosis with an excess of oxysterols such as 24-HC, 25-HC, 27-HC and 3 β-hydroxy-5-cholestenoic acid [10, 11]., which induces cytotoxicity [12–14]. However, the role of CYP7B1 expression with the development of NASH, the mechanisms underlying the downregulation of Cyp7b1 expression during NASH onset, and the consequences of this downregulation remain unclear.

Since two different NAFLD models show downregulation of *Cyp7b1* expression, we hypothesize that the hepatic expression of CYP7B1 may impact NASH development. In this study, we examined the expression of CYP7B1 and its roles on the inflammation and fibrosis induced in NASH. First, we demonstrate that hepatic CYP7B1 was significantly decreased in NASH model mice induced by methionine and choline-deficient (MCD) diet. This downregulation was already induced in the early stage of NASH development, and the adeno-associated virus (AAV)-mediated restoration of CYP7B1 expression reduced the expression of inflammation- and fibrosis-related genes in MCD diet-fed mice. In addition, accumulation of palmitic acid and induction of oxidative stress, suppressed the transcription of *Cyp7b1* in hepatoma cells.

## Materials and methods

### Materials

The following were purchased for this study: monoclonal rabbit anti-CYP7B1 (ab138497) from Abcam (Cambridge, United Kingdom); polyclonal rabbit anti-GAPDH (10494-1-AP) from Proteintech (Rosemont, IL); monoclonal rabbit anti-α-SMA (D4K9N) antibodies from Cell Signaling Technology (Danvers, MA); monoclonal mouse anti-ß-ACTIN (AC-15) from Sigma-Aldrich; and test kits for triglyceride (TG) E, cholesterol E, and transaminase CII from Wako.

### Animals and diets

Seven-week-old male C57BL/6J mice were purchased from CLEA Japan Inc., Tokyo, Japan. They were individually housed in a controlled environment (12-h light/dark cycle, lights-off between 9 PM and 9 AM, 25□, and controlled humidity). Distilled water was available ad libitum. All mice were acclimated to the control (Ctrl) diet for 1 week and then randomly fed with a methionine choline-deficient (MCD) diet or Ctrl diet for 8 weeks.

The mice were randomly assigned to one of the four experimental groups. Two of the four groups were intraperitoneally (i.p.) injected with AAV2/8-EGFP (mock) and the remaining two groups were injected with AAV2/8-FLAG-mCyp7b1 at 1 × 10^11^ virus genome per mouse. After injection, all mice were acclimated to the Ctrl diet for 1 week. Then, two groups from each AAV type were given free access to the Ctrl diet, while the remaining two groups to the MCD diet for 8 weeks.

To obtain liver and serum samples, the mice were euthanized under isoflurane anesthesia at the end of the experiment. The samples were stored at −80□ until further processing.

All animal experiments were performed according to the Guideline for the Care and Use of Laboratory Animals of the University of Tokyo and approved by the Animal Usage Committee of The University of Tokyo.

### Plasmid construction

PCR with appropriate primer sets used in this experiment are listed in Table S1. The DNA fragment encoding *Cyp7b1* (NM_007825) was amplified and cloned into p3×FLAG-CMV-10 vector (Sigma) using *Eco*RI/*Kpn*I restriction enzyme sites to construct pCMV10-3×FLAG-mCyp7b1.

Using pCMV10-3×FLAG-mCyp7b1 as a template, the coding sequence of the NH2-terminal FLAG-tagged mouse Cyp7b1 gene was also amplified and cloned into pAM.LSP1-EGFP using *Xba*I/*Eco*RV restriction enzyme sites to construct pAM.LSP1-3×FLAG-mCyp7b1.

### Adeno-associated virus preparation

The DNA fragment encoding N-terminal FLAG-tagged *Cyp7b1* was amplified and cloned into pAM.LSP1-EGFP (Children’s Medical Research Institute, Australia) using *Xba*I and *Eco*RV restriction enzyme sites to construct pAM.LSP1-FLAG-Cyp7b1.

On day 0, HEK293T cells were seeded in 150 mm dishes at 7 × 10^6^ for virus packaging. On day 1, the cells were transfected with FLAG-Cyp7b1 or empty adeno-associated vector along with AAV capsid 8 plasmid p5E18-VD2/8 (Penn Vector core, PA, USA) and helper plasmid pXX6-80 (NGVB, IN, USA) by polyethyleneimine (PEI)-max method. On day 3, the cells were switched to the medium supplemented with 2% (v/v) FBS and were harvested 60 h after transfection by scraping the medium. After centrifugation at 1,500 rpm for 5 min, the supernatant was discarded, and all pellets were resuspended by resuspension buffer (10 mM Tris, pH 8.0, 100 mM NaCl, 2 mM MgCl_2_) prior to three times freeze/thawing cycles. Centrifugation was conducted at 4,000 rpm for 15 min. Then, 50 U/mL benzonase was added to the supernatant before incubation at 37°C for 30 min; centrifugation was conducted again at 4,000 rpm for 10 min. The supernatants were subjected to CsCl density gradient fractionation. The peak fractions determined by the concentration of virus genome copies were collected. After dialysis with PBS (+) containing 5% (v/v) glycerol, RT-qPCR method was performed to determine the genome copy titers of AAV vectors using probes and primers targeting woodchuck posttranscriptional regulatory element region.

### Hepatic lipid extraction

Liver tissue was homogenized with 4 mL of chloroform/methanol (2:1, v/v) solution in a glass tube using a handy micro-homogenizer (Microtec, Chiba, Japan). To extract the lipids, the sample was vortexed and incubated on ice for 30 min. Afterward, for extraction, 1 mL of 50 mM NaCl was added and vortexed for 10 s before centrifugation at 1,500 g, 4□ for 30 min. An aspirator was used to remove the upper aqueous phase. The lipid-containing organic phase was collected in a new glass tube, washed twice with 1 mL of 0.36 M CaCl_2_/methanol (1:1, v/v), and then centrifuged at 1,500 g, 4□ for 10 min. The lower organic phase was collected in a measuring flask and filled up to 5 mL with chloroform. The extracted hepatic lipid was transferred to a new glass tube, air-replaced by nitrogen gas, and stored at –20□ until further examination.

### Hepatic and serum biochemistry

To determine the levels of hepatic cholesterol and TG, serum aspartate aminotransferase (AST), and alanine aminotransferase (ALT), kits purchased from FUJIFILM Wako Pure Chemical were used.

### Cell culture and cell treatment

Hepa1-6 cells (obtained from RIKEN Cell Bank, Japan) were maintained in complete DMEM supplemented with 10% (v/v) FBS, 100 unit/mL penicillin, and 100 μg/mL streptomycin. All cell cultures were maintained in a humidified incubator at 37°C under 5% CO_2_. On day 0, Hepa1-6 cells were seeded in a 6-well plate at 3.0 × 10^5^ cells/well or in 12-well plates at 2.0 × 10^5^ cells/well. On day 1, the cells were exposed to 0–400 μM oleic acid for 24 h, 0–800 μM palmitic acid for 15 h, or 0–200 μM H_2_O_2_ for 6 h. Subsequently, the cells were harvested for real-time (RT) quantitative PCR.

### Immunoblot analysis

Cells were lysed in lysis buffer (50 mM Tris□HCl, pH 8.0, 1 mM EDTA, 150 mM NaCl, 1% NP-40, and 0.25% sodium deoxycholate). To remove cellular debris, the lysate was centrifuged at 18,000 rpm for 10 min. The supernatants were adjusted to appropriate concentrations and treated with 6× Laemmli sample buffer (1 mM Tris□HCl, pH 6.8, 30% glycerol, 10% SDS, 600 mM dithiothreitol, and 0.03% bromophenol blue). To analyze the protein samples, SDS-PAGE was performed followed by immunoblot analysis with indicated antibodies. The samples were visualized using a western blotting detection reagent (Amersham Bioscience ECL, GE Healthcare Life Sciences) or Immobilon Western chemiluminescent HRP substrate (Merck Millipore).

### Real-time quantitative PCR

The total RNA from mouse liver or Hepa1-6 cells was extracted using ISOGEN (Nippon Gene) and reverse-transcribed using High-Capacity cDNA Reverse Transcription Kit (Thermo Fisher Scientific) following the manufacturer’s protocol. RT quantitative PCR was performed using the Applied Biosystems StepOnePlus™ RT PCR system (Thermo Fisher Scientific) with FastStart Universal SYBR Green Master (Roche Applied Science) following the manufacturer’s protocol. All relative mRNA expression levels were normalized with *18S*. Primer sequences are listed in Table S2.

### Statistical analysis

Data are presented as mean ± SD. Statistical analysis was performed using GraphPad Prism 9 (GraphPad Software). Significance through one-way analysis of variance with Tukey’s post hoc or the Student *t*-test was determined.

## Results

### Liver Cyp7b1 expression was decreased in MCD diet-induced NASH

From two data sets of micro-arrays, 45,101 genes demonstrated changes in leptin-deficient (*ob*/*ob*) mice and MCD diet-fed mice. *Cyp7b1* is predicted as one of the genes that exhibit a marked downregulation under the progression of NAFLD. To determine whether this downregulation attenuate the protein expression of Cyp7b1 in the progression of NASH, the mice were fed an MCD diet for 8 weeks. Compared with Ctrl diet-fed mice, MCD diet-fed mice demonstrated an increase in hepatic TG levels with the increase in inflammatory markers, serum aspartate aminotransferase (AST), and alanine aminotransferase (ALT) levels (Fig. 1A and B). Likewise, their hepatic expression of inflammatory cytokine-related genes, such as *Tnf-*α, *Il-1*β, and *Tlr4*, was significantly elevated. Moreover, markers for ER stress, C/EBP homologous protein (*Chop*), and activating transcription factor 4 (*Atf4*) were elevated in addition to the splicing of X-box binding protein 1 (*Xbp1*) mRNA. Fibrotic genes such as α*-Sma*, *Col1a1*, and *Tgf-*β significantly increased (Fig. 1C). Similar to gene expression, the protein expression of α-SMA in MCD diet-fed mouse liver was increased in 8 weeks (Fig. 1D). Thus, NASH was developed in 8 weeks of MCD diet feeding. Under this condition, Cyp7b1 expression level was downregulated in MCD diet fed mouse liver (Fig. 1E).

**Figure 1.**
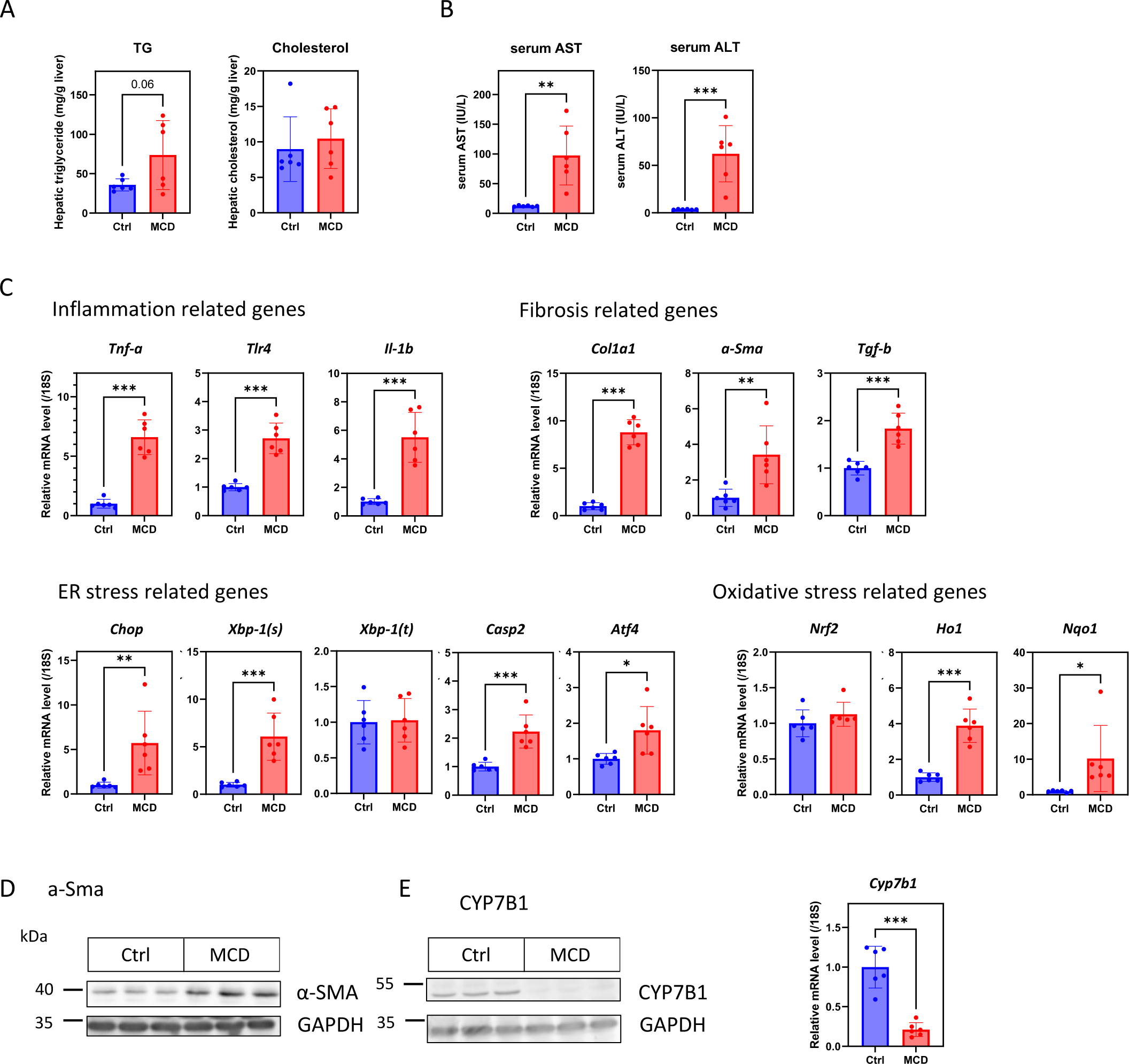
The mRNA and protein expression of CYP7B1 was decreased in NASH-developed mouse liver. C57BL/6J male mice were fed with MCD or Ctrl diet for 8 weeks. (A) Hepatic triglyceride and cholesterol levels. (B) Serum AST and serum ALT activity. (C) The mRNA of inflammation-related genes, fibrosis-related genes, ER stress-related genes, and oxidative stress-related genes. (D)The protein expression level of a-Sma. (E) The left panel shows CYP7B1 protein expression. The right panel shows Cyp7b1 mRNA expression. The data show mean ± SD. *p < 0.05, **p < 0.01, and ***p < 0.001.

### Restoration of CYP7B1 modestly ameliorates the progression of NASH by MCD diet

Figure 1 shows the downregulation of *Cyp7b1* induced in NASH-developed liver of MCD diet-fed mice presenting with lipid accumulation, inflammation, and fibrosis. To implicate CYP7B1 and NASH progression, the mice that had been fed Ctrl or MCD diet were induced with FLAG-tagged CYP7B1 using an AAV, referred to as AAV-FLAG-CYP7B1. Then, the effect of CYP7B1 expression on pathological features of NASH was analyzed by measuring the serum level of inflammatory-related enzymes and hepatic gene expressions. The endogenous CYP7B1 expression was decreased in MCD diet-fed mice infected with mock vectors or AAV-FLAG-CYP7B1. By intraperitoneal administration of AAV-FLAG-CYP7B1, the expression levels of CYP7B1 were similar in the MCD diet-fed mice and Ctrl diet-fed mice (Fig. 2A). An enforced expression of CYP7B1 significantly inhibited the upregulation of serum AST and ALT levels (Fig. 2B). Furthermore, among the inflammation-related genes and fibrosis markers that are elevated with the progression of NASH, *Tnf-a* and *Il-1b* in inflammatory genes, α*-Sma* and *Tgf-b* in fibrosis-related genes, and α-SMA protein expression in the liver were upregulated and resolved by CYP7B1 expression (Fig. 4C and D). Additionally, compared with mock-infected mice, MCD diet-fed mice demonstrated moderately decreased hepatic TG levels due to the restoration of CYP7B1 (Fig. 2E). These results suggest that the restored level of CYP7B1 attenuated hepatic inflammation and fibrosis.

**Figure 2.**
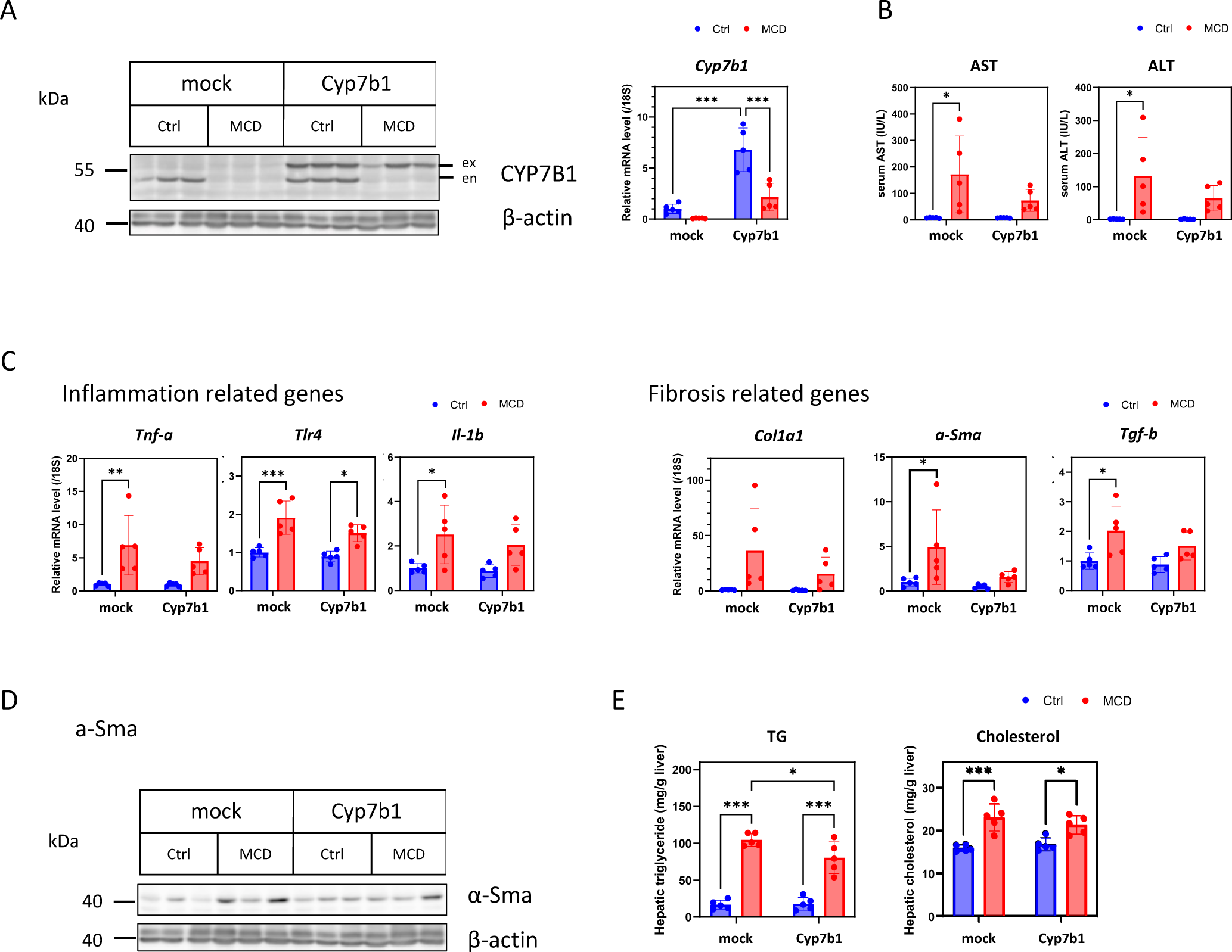
Effects of hepatic CYP7B1 restoration on the progression of NASH induced by MCD diet. C57BL/6J male mice injected by Mock or CYP7B1 expressing AAV-vector were fed with MCD or a Ctrl diet for 8 weeks. (A) The left panel shows CYP7B1 protein expression. The right panel shows Cyp7b1 mRNA expression. ex, exogenous; en, endogenous. (B) Serum AST and ALT activity. (C) The mRNA of Inflammation-related genes and fibrosis-related genes. (D) a-Sma expression. (E) Hepatic triglyceride and cholesterol. The right panel shows Cyp7b1 mRNA expression. The data show mean ± SD. *p < 0.05, **p < 0.01, and ***p < 0.001.

### Cyp7b1 expression was decreased in an early stage of NASH by lipotoxicity

To elucidate the mechanism of the downregulation of *Cyp7b1* in the early stage of NASH caused by the MCD diet, the feeding period was shortened to 1 week. As shown in Fig. 3A, 1-week MCD diet feeding caused a fivefold decrease in hepatic *Cyp7b1* mRNA level and reduced protein expression. To reveal the cause of *Cyp7b1* downregulation by MCD diet feeding, features and genes representing markers for the pathogenesis of NASH were examined. After 1 week of feeding, hepatic TG and total cholesterol levels increased (Fig. 3B), indicating the induction of lipid accumulation from the time point. In serum, increased circulating AST and ALT levels indicated hepatic injury (Fig. 3C). However, among inflammatory markers, only hepatic *Tnf-*α was significantly upregulated (Fig. 3D and S1A), as well as oxidative stress markers, represented in *Ho-1* and *Nqo-1* mRNA expression, after 1 week of MCD diet feeding (Fig. 3E). In contrast, among ER stress markers, *Chop* was the only one that increased (Fig. S1C), suggesting that only some inflammatory pathways and stress signals were induced in this stage of steatosis.

**Figure 3.**
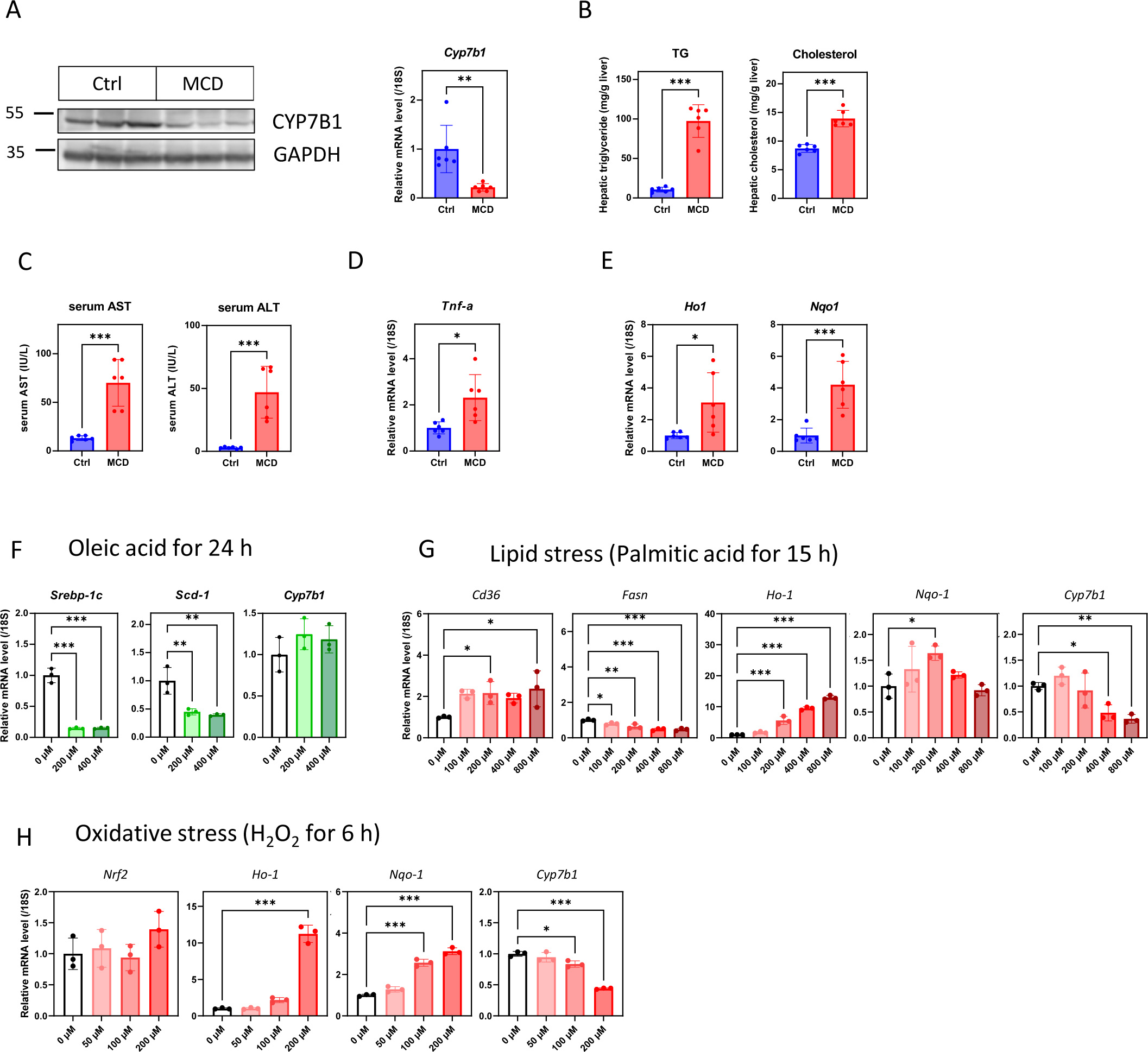
Lipotoxicity in the early stage of NASH suppresses the expression of CYP7B1. C57BL/6J male mice were fed with MCD or Ctrl diet for 1 week (A–E). (A) The left panel shows CYP7B1 protein expression, while the right panel shows Cyp7b1 mRNA expression. (B) Hepatic triglyceride and cholesterol levels. (C) Serum AST and serum ALT activity. (D, E) The hepatic mRNA levels of Tnf-a (D) and oxidative stress-related genes (E). (F) Hepa1-6 cells were exposed to 0–400 μM oleic acid for 24 h. Ethanol (conjugated with BSA in PBS) served as a vehicle. (G) Hepa1-6 cells were exposed to 0–800 μM palmitic acid for 15 h. Ethanol (conjugated with BSA in PBS) served as a vehicle. (H) Hepa1-6 cells were treated with 0–200 μM H_2_O_2_ for 6 h. PBS was used as a vehicle. The data show mean ± SD. *p < 0.05, **p < 0.01, and ***p < 0.001.

To confirm the progression of NASH during *Cyp7b1* downregulation, features and genes related to a NASH pathological hit were measured. NAFLD is defined by the excessive accumulation of lipids within hepatocytes. Since hepatic lipid accumulation was observed even in week 1 of MCD diet feeding (Fig. 3B), *Cyp7b1* expression might have been affected by the accumulation of fatty acids. To verify this hypothesis, murine hepatoma Hepa1-6 cells were treated with indicated concentrations of oleic acid and palmitic acid. The results revealed that the expression of *Cyp7b1* gene was not affected by the exposure to oleic acid, whereas positive controls, *Srebp-1c* and *Scd-1*, exhibited 5- and 2-fold decreases, respectively (Fig. 3F). In contrast, palmitic acid reduced the same expression in a dose-dependent manner, beginning with 400 µM. Exposure to palmitic acid upregulated the gene expressions of cluster of differentiation 36 (*Cd36*), which increase by the presence of the fatty acid, and downregulated fatty acid synthase (*Fasn*), which is decreased by its product. Moreover, the expression of the *Ho-1* gene, a marker of oxidative stress, was also increased in a dose-dependent manner (Fig. 3G). These results indicate that the downregulation of *Cyp7b1* might be a consequence of lipotoxicity from accumulating free fatty acids, specifically saturated fatty acids.

To confirm the effect of oxidative stress on *Cyp7b1* expression, Hepa1-6 cells were exposed to various concentrations of H_2_O_2_ for 6 h. The expressions of *Ho-1* and *Nqo-1* were significantly upregulated with 200 µM H_2_O_2_ and >100 µM H_2_O_2_, respectively (Fig. 3H). These results indicate that H_2_O_2_ treatment induces oxidative stress in Hepa1-6 cells. Gene expression of *Cyp7b1* was significantly decreased by 100 µM H_2_O_2_, and its expression was reduced to approximately 50% by 200 µM H_2_O_2_.

## Discussion

NASH is a progressive stage in NAFLD, characterized by ballooning degeneration, inflammation, and fibrosis. Its pathogenesis is multifactorial, but the link between lipid accumulation and inflammation remains unclear. In the present study, we showed that CYP7B1 partially related to NASH progression and its gene expression was regulated by the accumulation of fatty acid.

The gene expression profiles in GSE35961 are stored in the Gene Expression Omnibus database (National Institutes of Health, Bethesda, MD). The expression of hepatic *Cyp7b1* mRNA was significantly downregulated in MCD plus high-fat diet mice compared with that in normal diet-fed mice, similar to results for GSE49195 in which *Cyp7b1* gene expression in the fatty liver of obese ob/ob mice was lower than that of wild-type mice [7, 8]. Since both two data sets from different NAFLD models, dietary and genetically induced, showed the downregulation of the *Cyp7b1* gene expression, the pathology of NAFLD including NASH might be related to *Cyp7b1*. Not only gene expression but also protein expression of CYP7B1 was strongly downregulated in NASH-progressed liver by MCD diet feeding (Fig. 1). Because 25-hydroxycholesterol and 27-hydroxycholesterol, the most well-known substrate of CYP7B1, are toxic oxysterols, the decrease in CYP7B1 protein expression might cause liver injury due to accumulation [13–15].

Of note, gene transfer mediated by AAV vectors shows long-term and persistent transgene expression with a mild level of immune response [16, 17]. In the present study, AAV seemed a suitable system to deliver Cyp7b1 gene expression during 8 weeks of MCD diet feeding. The restoration of CYP7B1 by AAV delivery transgene alleviated inflammation and fibrosis in MCD diet-induced NASH. CYP7B1 was downregulated in MCD diet-fed animals showing NASH progression, but its restoration inhibited NASH progression, therefore indicating that decreased CYP7B1 expression enhances NASH progression. Our results support non-NAFLD studies reporting genetic deficiency of *CYP7B1* that resulted in hepatic inflammation and progression to severe fibrosis in the neonatal stage [18, 19]. Furthermore, Kakiyama et al. reported that the liver of mice with NASH that received western diet feeding also showed a decrease of *Cyp7b1* and that dietary coffee restored the expression of *Cyp7b1*, thereby moderated liver injury [20].

The induction of CYP7B1 downregulation was rapid and constant, compared to other pathological features such as some of the inflammatory cytokines and fibrosis-related genes that did not change after 1 week of MCD diet feeding. This suggests that the downregulation of CYP7B1 is rather an upstream than a downstream in the progression of chronic liver injury. However, some of the “hits” for NASH progression were induced, such as accumulation of hepatic TG or increase in proinflammatory genes and oxidative stress-related genes in the early stage of NASH after 1 week of MCD diet feeding.

Lipid accumulation was induced in the liver of mice since week 1 of MCD diet feeding. Among fatty acids, oleic acid and palmitic acid are the most abundant circulating fatty acids in NASH patients and significantly increase in healthy subjects [21]. Downregulation of *Scd-1* and *Srebp-1c* in oleic acid-treated cells confirmed that the administration of oleic acid disturbed intracellular lipid metabolism [22]. Likewise, the upregulation of *Cd36* and downregulation of *Fasn* in palmitic acid-treated cells indicated the regulatory function since palmitic acid increases the gene expression of fatty acid transporter and suppresses its synthetic enzyme via negative feedback. Palmitic acid at high concentrations (over 400 μM) downregulated *Cyp7b1*, while oleic acid induced no change (Fig. 3F and G). This indicates that a saturated fatty acid known for its apoptotic, lipotoxic, and oxidative stress-inducing features may have caused Cyp7b1 downregulation [23–25]. *Cyp7b1* expression was downregulated with the increase in oxidative stress markers.

Another hit induced in 1-week MCD diet feeding is oxidative stress. Together with this, 6 h of H_2_O_2_ treatment in Hepa1-6 cells, which upregulated oxidative stress markers, also exhibited a remarkable decrease in Cyp7b1 expression. This result supports a study of bile duct ligation-induced oxidative stress and liver injury, in which the hepatic *Cyp7b1* expression diminished after bile duct ligation and further dropped when *Nrf2*, a nuclear factor regulating the expression of antioxidation enzymes, was ablated [26]. Furthermore, depending on the *Cyp7b1* downregulation after palmitic acid exposure, oxidative markers in palmitic acid-treated cells were increased. Taking these together, oxidative stress may also cause the downregulation of *Cyp7b1*.

In conclusion, our findings suggest a novel possibility of the implication of CYP7B1 in the progression of NASH. The lipotoxicity caused by an accumulation of saturated fatty acids suppressed *Cyp7b1* expression in the early stage of NAFLD liver. The loss of CYP7B1 aggravates inflammatory and fibrosis markers that were ameliorated by the restoration of CYP7B1 expression, suggesting the downregulation of *Cyp7b1* as a “hit” in the NASH pathogenesis. Further studies are needed to evaluate CYP7B1 as a therapeutic target for patients with NASH.

## Data availability

The data that support the findings of this study are available from the corresponding author upon reasonable request.

## Author contributions

**Yuichi Watanabe:** Writing - Original Draft, Visualization, Supervision, Funding acquisition **Kyouhei Kinoshita:** Visualization, Investigation, Validation **Napatsorn Dumrongkulchart:** Visualization, Investigation, Validation **Takashi Sasaki:** Investigation, Validation **Makoto Shimizu:** Investigation, Validation **Yoshio Yamauchi:** Investigation, Validation **Ryuichiro Sato:** Conceptualization, Writing - Review & Editing, Project administration, Funding acquisition

## Acknowledgments

We thank Prof. I. Alexander (Children’s Medical Research Institute, Australia) for providing pAM-LSP1-EGFP plasmid. We thank National gene vector biorepository (University of North Carolina at Chapel Hill, USA) for pXX6-80 plasmid, a helper vector used for the package of CYP7B1-expressing AAV8, and Penn Vector Core (University of Pennsylvania, USA) for p5E18-VD2/8 plasmid. We also thank Dr. Tohru Itoh (Institute for Quantitative Biosciences, The University of Tokyo) for kindly teaching AAV production and providing instruments. This work was supported by Japan Society for the Promotion of Science (JSPS) KAKENHI Grant-in-Aid for Scientific Research (S) JP15H05781 and (A) JP20H00408 (to R. S.), Grant-in-Aid for Young Scientists JP19K15764 (to Y. W.), and the Japan Agency for Medical Research and Development (AMED-CREST) Grant 16gm0910008h0001 (to R. S.). The authors would like to thank Enago (www.enago.jp) for the English language review.

---

**Figure S1.**
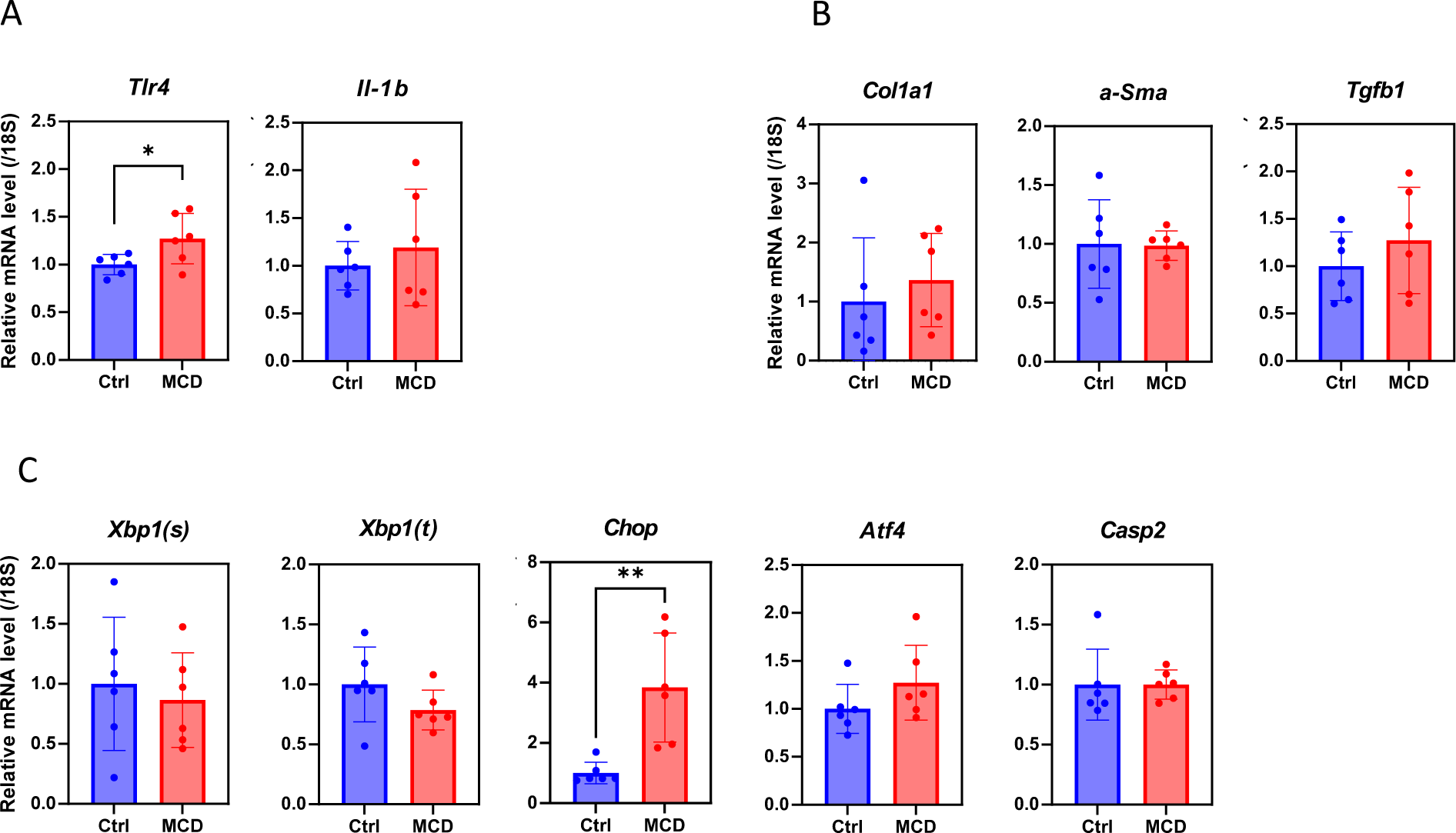
Hepatic mRNA levels in C57BL/6J male mice that were fed with MCD or Ctrl diet for 1 week. (A) Inflammation-related genes. (B) Fibrosis-related genes. (C) ER stress-related genes. The data show mean ± SD. *p < 0.05, **p < 0.01.

## Reference

[1] Z.M. Younossi, A.B. Koenig, D. Abdelatif, Y. Fazel, L. Henry, M. Wymer, Global epidemiology of nonalcoholic fatty liver disease-Meta-analytic assessment of prevalence, incidence, and outcomes, Hepatology, 64 (2016) 73–84.

[2] H. Tilg, A.R. Moschen, Evolution of inflammation in nonalcoholic fatty liver disease: the multiple parallel hits hypothesis, Hepatology, 52 (2010) 1836–1846.

[3] J.V. Lazarus, H.E. Mark, Q.M. Anstee, J.P. Arab, R.L. Batterham, L. Castera, H. Cortez-Pinto, J. Crespo, K. Cusi, M.A. Dirac, S. Francque, J. George, H. Hagstrom, T.T. Huang, M.H. Ismail, A. Kautz, S.K. Sarin, R. Loomba, V. Miller, P.N. Newsome, M. Ninburg, P. Ocama, V. Ratziu, M. Rinella, D. Romero, M. Romero-Gomez, J.M. Schattenberg, E.A. Tsochatzis, L. Valenti, V.W. Wong, Y. Yilmaz, Z.M. Younossi, S. Zelber-Sagi, N.C. Consortium, Advancing the global public health agenda for NAFLD: a consensus statement, Nat Rev Gastroenterol Hepatol, 19 (2022) 60–78.

[4] A.R. Stiles, J.G. McDonald, D.R. Bauman, D.W. Russell, CYP7B1: one cytochrome P450, two human genetic diseases, and multiple physiological functions, J Biol Chem, 284 (2009) 28485–28489.

[5] J.Y. Chiang, Regulation of bile acid synthesis: pathways, nuclear receptors, and mechanisms, J Hepatol, 40 (2004) 539–551.

[6] A.V. Yantsevich, Y.V. Dichenko, F. Mackenzie, D.V. Mukha, A.V. Baranovsky, A.A. Gilep, S.A. Usanov, N.V. Strushkevich, Human steroid and oxysterol 7alpha-hydroxylase CYP7B1: substrate specificity, azole binding and misfolding of clinically relevant mutants, FEBS J, 281 (2014) 1700–1713.

[7] S. Antherieu, D. Le Guillou, C. Coulouarn, K. Begriche, V. Trak-Smayra, S. Martinais, M. Porceddu, M.A. Robin, B. Fromenty, Chronic exposure to low doses of pharmaceuticals disturbs the hepatic expression of circadian genes in lean and obese mice, Toxicol Appl Pharmacol, 276 (2014) 63–72.

[8] Y. Kita, T. Takamura, H. Misu, T. Ota, S. Kurita, Y. Takeshita, M. Uno, N. Matsuzawa-Nagata, K. Kato, H. Ando, A. Fujimura, K. Hayashi, T. Kimura, Y. Ni, T. Otoda, K. Miyamoto, Y. Zen, Y. Nakanuma, S. Kaneko, Metformin prevents and reverses inflammation in a non-diabetic mouse model of nonalcoholic steatohepatitis, PLoS One, 7 (2012) e43056.

[9] G. Kakiyama, D. Marques, R. Martin, H. Takei, D. Rodriguez-Agudo, S.A. LaSalle, T. Hashiguchi, X. Liu, R. Green, S. Erickson, G. Gil, M. Fuchs, M. Suzuki, T. Murai, H. Nittono, P.B. Hylemon, H. Zhou, W.M. Pandak, Insulin resistance dysregulates CYP7B1 leading to oxysterol accumulation: a pathway for NAFL to NASH transition, J Lipid Res, 61 (2020) 1629–1644.

[10] I. Ueki, A. Kimura, A. Nishiyori, H.L. Chen, H. Takei, H. Nittono, T. Kurosawa, Neonatal cholestatic liver disease in an Asian patient with a homozygous mutation in the oxysterol 7alpha-hydroxylase gene, Journal of pediatric gastroenterology and nutrition, 46 (2008) 465–469.

[11] K.D. Setchell, M. Schwarz, N.C. O’Connell, E.G. Lund, D.L. Davis, R. Lathe, H.R. Thompson, R. Weslie Tyson, R.J. Sokol, D.W. Russell, Identification of a new inborn error in bile acid synthesis: mutation of the oxysterol 7alpha-hydroxylase gene causes severe neonatal liver disease, J Clin Invest, 102 (1998) 1690–1703.

[12] Y. Urano, D.K. Ho Vo, A. Hirofumi, N. Noguchi, 24(S)-Hydroxycholesterol induces ER dysfunction-mediated unconventional cell death, Cell Death Discov, 5 (2019) 113.

[13] A. Asghari, T. Ishikawa, S. Hiramitsu, W.R. Lee, J. Umetani, L. Bui, K.S. Korach, M. Umetani, 27-Hydroxycholesterol Promotes Adiposity and Mimics Adipogenic Diet-Induced Inflammatory Signaling, Endocrinology, 160 (2019) 2485–2494.

[14] Y. Watanabe, T. Sasaki, S. Miyoshi, M. Shimizu, Y. Yamauchi, R. Sato, Insulin-induced genes INSIG1 and INSIG2 mediate oxysterol-dependent activation of the PERK-eIF2alpha-ATF4 axis, J Biol Chem, 297 (2021) 100989.

[15] Y. Yamanashi, T. Takada, Y. Tanaka, Y. Ogata, Y. Toyoda, S.M. Ito, M. Kitani, N. Oshida, K. Okada, J. Shoda, H. Suzuki, Hepatic Niemann-Pick C1-Like 1 exacerbates non-alcoholic fatty liver disease by re-absorbing specific biliary oxysterols, Biomed Pharmacother, 156 (2022) 113877.

[16] S.C. Cunningham, A.P. Dane, A. Spinoulas, I.E. Alexander, Gene Delivery to the Juvenile Mouse Liver Using AAV2/8 Vectors, Molecular Therapy, 16 (2008) 1081–1088.

[17] A.P. Dane, S.J. Wowro, S.C. Cunningham, I.E. Alexander, Comparison of gene transfer to the murine liver following intraperitoneal and intraportal delivery of hepatotropic AAV pseudo-serotypes, Gene Ther, 20 (2013) 460–464.

[18] D. Dai, P.B. Mills, E. Footitt, P. Gissen, P. McClean, J. Stahlschmidt, I. Coupry, J. Lavie, F. Mochel, C. Goizet, T. Mizuochi, A. Kimura, H. Nittono, K. Schwarz, P.J. Crick, Y. Wang, W.J. Griffiths, P.T. Clayton, Liver disease in infancy caused by oxysterol 7 alpha-hydroxylase deficiency: successful treatment with chenodeoxycholic acid, J Inherit Metab Dis, 37 (2014) 851–861.

[19] J.Y. Chen, J.F. Wu, A. Kimura, H. Nittono, B.Y. Liou, C.S. Lee, H.S. Chen, Y.C. Chiu, Y.H. Ni, S.S. Peng, W.T. Lee, I.J. Tsai, M.H. Chang, H.L. Chen, AKR1D1 and CYP7B1 mutations in patients with inborn errors of bile acid metabolism: Possibly underdiagnosed diseases, Pediatr Neonatol, 61 (2020) 75–83.

[20] G. Kakiyama, K. Minowa, D. Rodriguez-Agudo, R. Martin, H. Takei, K. Mitamura, S. Ikegawa, M. Suzuki, H. Nittono, M. Fuchs, D.M. Heuman, H. Zhou, W.M. Pandak, Coffee modulates insulin-hepatocyte nuclear factor-4alpha-Cyp7b1 pathway and reduces oxysterol-driven liver toxicity in a nonalcoholic fatty liver disease mouse model, Am J Physiol Gastrointest Liver Physiol, 323 (2022) G488–G500.

[21] I. Tavares De Almeida, H. Cortez-Pinto, G. Fidalgo, D. Rodrigues, M.E. Camilo, Plasma total and free fatty acids composition in human non-alcoholic steatohepatitis, Clinical Nutrition, 21 (2002) 219–223.

[22] J. Ou, H. Tu, B. Shan, A. Luk, R.A. DeBose-Boyd, Y. Bashmakov, J.L. Goldstein, M.S. Brown, Unsaturated fatty acids inhibit transcription of the sterol regulatory element-binding protein-1c (SREBP-1c) gene by antagonizing ligand-dependent activation of the LXR, Proc Natl Acad Sci U S A, 98 (2001) 6027–6032.

[23] Y.S. Lee, S.Y. Kim, E. Ko, J.H. Lee, H.S. Yi, Y.J. Yoo, J. Je, S.J. Suh, Y.K. Jung, J.H. Kim, Y.S. Seo, H.J. Yim, W.I. Jeong, J.E. Yeon, S.H. Um, K.S. Byun, Exosomes derived from palmitic acid-treated hepatocytes induce fibrotic activation of hepatic stellate cells, Sci Rep, 7 (2017) 3710.

[24] N.P. Zhang, X.J. Liu, L. Xie, X.Z. Shen, J. Wu, Impaired mitophagy triggers NLRP3 inflammasome activation during the progression from nonalcoholic fatty liver to nonalcoholic steatohepatitis, Lab Invest, 99 (2019) 749–763.

[25] Y. Ogawa, K. Imajo, Y. Honda, T. Kessoku, W. Tomeno, S. Kato, K. Fujita, M. Yoneda, S. Saito, Y. Saigusa, H. Hyogo, Y. Sumida, Y. Itoh, K. Eguchi, T. Yamanaka, K. Wada, A. Nakajima, Palmitate-induced lipotoxicity is crucial for the pathogenesis of nonalcoholic fatty liver disease in cooperation with gut-derived endotoxin, Sci Rep, 8 (2018) 11365.

[26] L.M. Aleksunes, A.L. Slitt, J.M. Maher, M.Z. Dieter, T.R. Knight, M. Goedken, N.J. Cherrington, J.Y. Chan, C.D. Klaassen, J.E. Manautou, Nuclear factor-E2-related factor 2 expression in liver is critical for induction of NAD(P)H:quinone oxidoreductase 1 during cholestasis, Cell Stress Chaperones, 11 (2006) 356–363.

