## Supplemental Tables for "Downregulation of CYP7B1 caused by lipotoxicity associates with the progression of non-alcoholic steatohepatitis"

**Table S1 Sequences of PCR primers for cloning in expression vectors**

| PCR primers for cloning | Sequences (5’ to 3’) |
| --- | --- |
| **pCMV10-3×FLAG-mCyp7b1** | |
| mouseCyp7b1-EcoRI-Fw | AAATTTATAGAATTCCCAGGGAGCCACGACCCTAGAT |
| mouseCyp7b1-KpnI-Rv | AAATTTATAGGTACCTCAGCTTCTCCAAGATTTTGCTTTG |
|  | underline shows *Eco*RI and *Kpn*I site, respectively |
| **pAM.LSP1-3×FLAG-mCyp7b1** | |
| FLAG-mCyp7b1-XbaI-Fw | TTTATATCTAGAATGGACTACAAAGACCATGACG |
| FLAG-mCyp7b1-EcoRV-Rv | TTTATAGATATCTCAGCTTCTCCAAGATTTTGC |
|  | underline shows *Xba*I and *Eco*RV site, respectively |

**Table S2 Sequences of primers for qPCR**

| Targets | Forward primers (5’ to 3’) | | Reverse primers (5’ to 3’) | |
| --- | --- | --- | --- | --- |
| **Mouse** | | | | |
| *18s* | | ACCGCAGCTAGGAATAATGGA | | GCCTCAGTTCCGAAAACCA |
| *Cyp7b1* | | TTCCTCCACTCATACACAATG | | CGTGCTTTTCTTCTTACCATC |
| *Tnf-α* | | CAGCCGATGGGTTGTACCTT | | GGCAGCCTTGTCCCTTGA |
| *Tlr4* | | GTTCTCTCATGGCCTCCACT | | ATTAGGAACTACCTCTATGCAGGG |
| *Il-1ß* | | AGTTGACGGACCCCAAAAGA | | GGACAGCCCAGGTCAAAGG |
| *Col1a1* | | TTCTCCTGGCAAAGACGGACTCAA | | AGGAAGCTGAAGTCATAACCGCCA |
| *α-Sma* | | CCACCGCAAATGCTTCTAAGT | | GGCAGGAATGATTTGGAAAGG |
| *Tgf-ß* | | CGCCGCAGGCTCCTT | | CGTGAACTGATTTGGATCTTTGC |
| *Chop* | | CTGCCTTTCACCTTGGAGAC | | CGTTTCCTGGGGATGAGATA |
| *Xbp-1 (s)* | | GAGTCCGCAGCAGGTG | | GTGTCAGAGTCCATGGGA |
| *Xbp-1 (t)* | | GAGCAGCAAGTGGTGGATTT | | CCGTGAGTTTTCTCCCGTAA |
| *Casp2* | | CAATGCTAACTGTCCAAGTCTA | | GGGGATTGTGTGTGGTTCTT |
| *Atf4* | | ATGGCCGGCTATGGATGAT | | CGAAGTCAAACTCTTTCAGATCCATT |
| *Nrf2* | | CCGCTACACCGACTACGATT | | ACCTTCATCACCAACCCAAG |
| *Ho1* | | GTGTCCAGAGAAGGCTTTAAGCT | | TCTGCTTGTTGCGCTCTATCT |
| *Nqo1* | | GAAGCTGCAGACCTGGTGAT | | GCTACGAGCACTCTCTCTAAACC |
| *Srebp-1c* | | GAGCCATGGATTGCACATTT | | CGGGAAGTCACTGTCTTGGT |
| *Scd-1* | | CCGGAGACCCCTTAGATCGA | | TAGCCTGTAAAAGATTTCTGCAAACC |
| *Cd36* | | CTTCCACATTTCCTACATGCAA | | ATCCAGTTATGGGTTCCACATC |
| *Fasn* | | TGCTCCCAGCTGCAGGC | | GCCCGGTAGCTCTGGGTGTA |
